# Temporal learning of bottom-up connections via spatially nonspecific top-down inputs

**DOI:** 10.1101/649798

**Authors:** Jung Hoon Lee, Mean-Hwan Kim, Sujith Vijayan

## Abstract

In the brain, high-order and low-order areas are connected via bottom-up connections (from low-order to high-order areas) and top-down connections (from high-order to low-order areas). While bottom-up signals are thought to be critical in generating perception, functions of top-down signals have not been clearly delineated. One popular theory is that top-down inputs modify the activity of specific cell assemblies to modulate responses to bottom-up inputs. However, a different line of studies proposes that not all top-down inputs are specifically delivered. As the leading theories cannot account for nonspecific top-down inputs, we seek potential functions of nonspecific top-down signals using network models in our study. Our simulation results suggest that top-down inputs can regulate low-order area responses by providing temporal information even without spatial specificity. Specifically, the temporal information in nonspecific top-down inputs can weaken the undesired bottom-up connections, contributing to bottom-up connections’ learning. Further, we found that cortical rhythms (synchronous oscillatory neural responses) are critical in the proposed learning process of bottom-up connections in our model.

## 1. Introduction

The brain is thought to have hierarchical structures [1, 2, 3, 4, 5, 6], in which high-order areas rely on low-order areas for their functions. For instance, the prefrontal cortex needs the outputs of the visual cortices to access visual information. This hierarchy of the brain has inspired deep neural networks (DNN) [7, 8], in which information propagates from input to output layers in a feedforward fashion. However, the brain is not just a group of feedforward networks. Instead, high- and low-order areas in the brain are reciprocally and intensively connected [3, 5, 9, 10], and a line of studies suggests that both bottom-up (i.e., feedforward), from low-order to high-order areas, and top-down (i.e., feedback) signals, from high-order to low-order areas, are critical in cognitive functions [11, 12, 13, 14]. Bottom-up signals have been considered responsible for sensory signal processing, but functions of top-down signals remain poorly understood.

A few leading theories propose potential functions of top-down signals. One theory is that top-down inputs mediate selective attention or endogenous contexts, allowing sensory areas to respond selectively to more behaviorally important stimuli [15, 16]. Specifically, it is proposed that top-down inputs promote responses of a selected population of homogeneous neurons (i.e., an assembly) to win the competition among cell assemblies in low-order sensory areas; see [17] for a review. A second theory is that top-down inputs mediate expectations according to predictive coding theory [18, 19]. These theories assume that top-down signals can target specific cell assemblies. However, a different line of studies suggests that some top-down signals may not be target-specific [20, 21, 22, 23, 24]. Without target specificity, they cannot mediate attention or expectation, as the earlier theories suggest, indicating that nonspecific top-down inputs could play a different role. In this study, we use network models to pursue their potential functions.

Even without spatial specificity/information, top-down signals can still mediate temporal information onto target assemblies and evoke disparate responses depending on their exact arrival times. Importantly, the time-dependent efficacy of spikes can be profoundly impacted when target neurons are in synchronous and oscillatory states such as PING rhythms (oscillations at a gamma frequency generated by the interplay between excitatory and inhibitory neurons); see [25]. If top-down signals arrive immediately after inhibitory neurons fire synchronously, they will hardly entrain target neurons due to strong inhibition. In contrast, if top-down inputs arrive right before a rhythm’s cycle, they will entrain target neurons quite reliably. Further, we note that coordinated rhythmic activity is thought to mediate interareal communications in the brain [26] and allow reliable learning via spike-time dependent plasticity (STDP) [27]. Inspired by this line of ideas, we hypothesize that temporal information encoded in nonspecific top-down signals can help bottom-up signals propagate effectively by modulating the strength of bottom-up connections.

To address this hypothesis, we construct a network model consisting of one low-order and one high-order area. Specifically, we consider two inhomogeneous cell assemblies in a low-order area and test the effects of nonspecific top-down signals on the evolution of connections from the two assemblies to a high-order area. Our simulation results suggest that 1) nonspecific top-down connections can suppress undesired bottom-up connections and 2) the power of gamma rhythms in lower-order areas is correlated with the change in connection strength of connections between lower-order areas to higher-order areas, which reflects the degree of learning. These results suggest that even without spatial specificity, temporal information mediated by top-down inputs can change the strength of bottom-up signals. Furthermore, we note that STDP connections are known to generate runaway neural activity [28], which prevents spiking neural networks (SNNs) from learning practical tasks. Since in our simulations, nonspecific top-down inputs suppress undesired connections, we propose that nonspecific top-down inputs can help SNNs learn by preventing runaway activity in them; see Discussion.

## 2. Methods

To seek potential roles of nonspecific top-down inputs in bottom-up connections’ learning, we used network models to study nonspecific top-down inputs’ contribution to the formulation of selectivity in bottom-up connections.

### 2.1 Network Model

Our network model consists of three cell assemblies (Fig. 1). Two assemblies (PA, NPA) represent a low-order area, and the other (HA) represents a high-order area. PA and NPA are different only in their external inputs mimicking sensory inputs (Fig. 1). Based on the observations [3, 9] that bottom-up connections mainly target the granular layer (L4) and that top-down connections avoid L4, we constructed each assembly with two layers, superficial layer (L2/3) and L4. In each layer, excitatory neurons interact with inhibitory neurons. Commonly, excitatory neurons are referred to as pyramidal neurons due to their shape, and most inhibitory neurons are known to express parvalbumin (PV). Thus, excitatory and inhibitory neurons are referred to as Pyr and PV neurons in this study. Each neuron type is connected to other neuron types in the same assembly. The connections are randomly established using connection probabilities determined by pre- and postsynaptic neuron types. The strengths of connections also depend on pre- and postsynaptic neuron types. The selected values for connection probabilities and strengths are listed in Table 1. The three assemblies in the model are connected with one another via inter-assembly (i.e., interareal) connections (Table 1). We used the peer-reviewed open-source simulation platform NEST [29] to build the network models. All parameters not specified here are taken from the default parameters of the NEST package [29].

**Figure 1:**
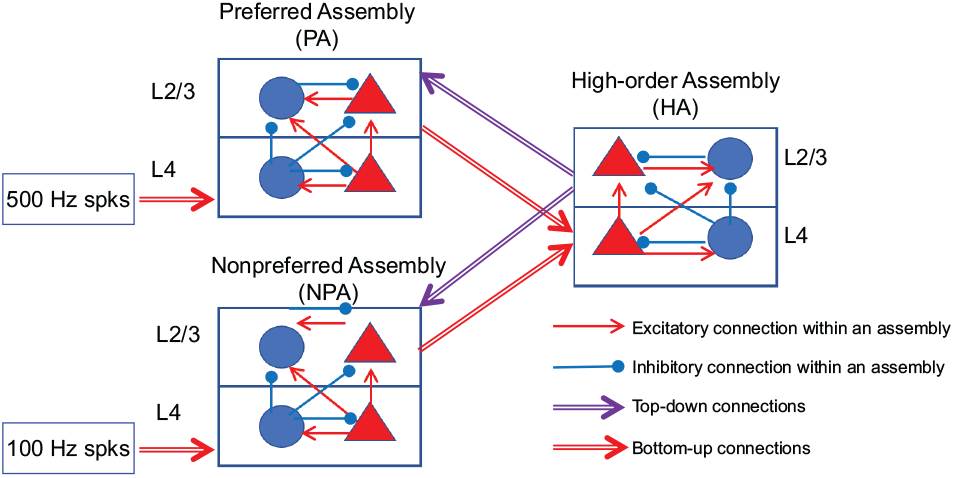
Schematics of the model. The model consists of three assemblies (PA, NPA, and HA). In superficial (L2/3) and granular (L4) layers, Pyr (red triangles) and PV (blue circles) interact with one another. PA and NPA together represent the low-order area, and they receive disparate external inputs. HA represents the high-order area. All three assemblies interact through layer-specific top-down and bottom-up connections.

**Table 1:**
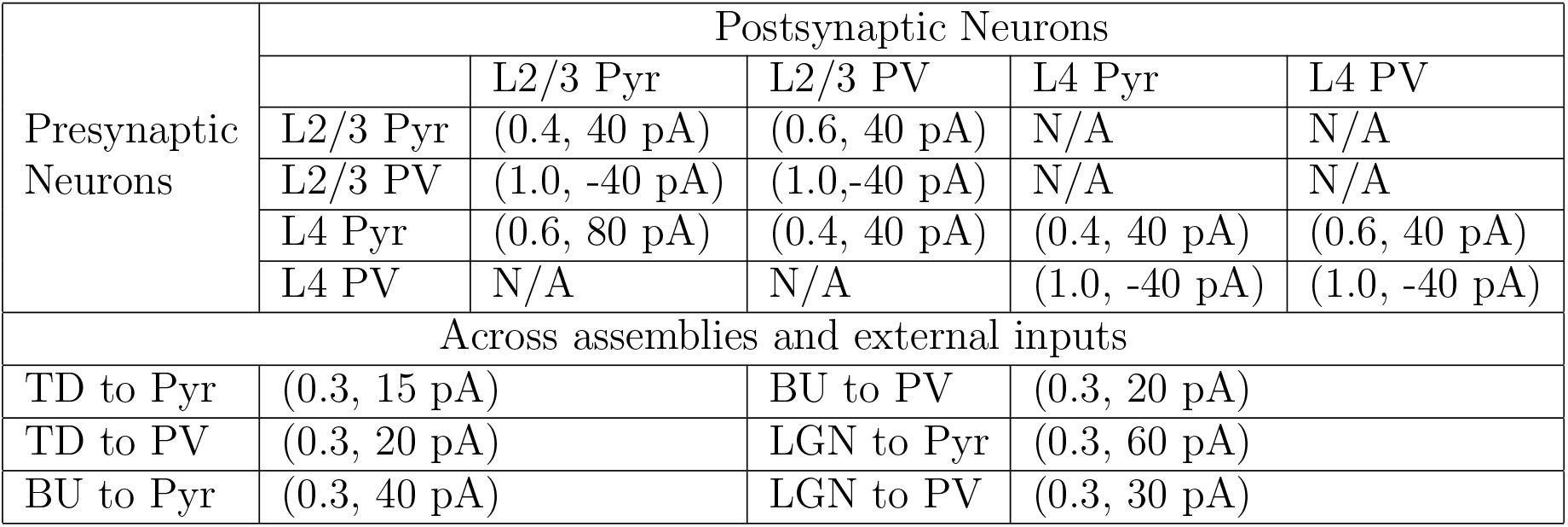
Connections in the network model. Below, the connection probability and strength of each connection type are shown in parentheses. TD, BU, and LGN represent top-down, bottom-up, and lateral geniculate nucleus connections, respectively. In the model, we consider both low-order and high-order areas. Only low-order areas, which model sensory cortices, receive LGN inputs. Additionally, Pyr and PB neurons receive 1050 Hz and 1000 Hz background inputs, respectively, via 100 pA connections.

| Presynaptic Neurons | Postsynaptic Neurons |  |  |  |  |
| --- | --- | --- | --- | --- | --- |
|  |  | L2/3 Pyr | L2/3 PV | L4 Pyr | L4 PV |
|  | L2/3 Pyr | (0.4, 40 pA) | (0.6, 40 pA) | N/A | N/A |
|  | L2/3 PV | (1.0, -40 pA) | (1.0, -40 pA) | N/A | N/A |
|  | L4 Pyr | (0.6, 80 pA) | (0.4, 40 pA) | (0.4, 40 pA) | (0.6, 40 pA) |
|  | L4 PV | N/A | N/A | (1.0, -40 pA) | (1.0, -40 pA) |
| Across assemblies and external inputs |  |  |  |  |  |
| TD to Pyr | (0.3, 15 pA) |  | BU to PV | (0.3, 20 pA) |  |
| TD to PV | (0.3, 20 pA) |  | LGN to Pyr | (0.3, 60 pA) |  |
| BU to Pyr | (0.3, 40 pA) |  | LGN to PV | (0.3, 30 pA) |  |

### 2.2 Neuron Model

In the model, all neurons are current-based leaky-integrate fire (LIF) neurons. The dynamics of subthreshold membrane potentials are described by Eq. 1.

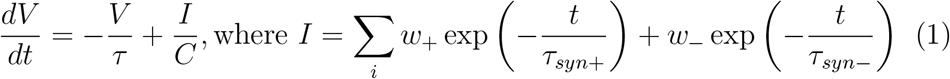

where *w*_+_ and *τ_syn_*_+_ represent the synaptic strengths and synaptic time scales of excitatory synapses and where *w_−_* and *τ_syn−_* represent the synaptic strengths and synaptic time scales of inhibitory synapses. The parameters used in this study are listed in Table 2.

**Table 2:** Parameters for neurons and synaptic inputs. Synaptic inputs decay at time scales (*τ_s_*) depending on the identities of presynaptic and postsynaptic neurons.

| Param | Value | Param | Value |
| --- | --- | --- | --- |
| Membrane constant | 10 ms | $\tau_s(Pyr \rightarrow PV)$ | 2 ms |
| Spike threshold | -50 mV | $\tau_s(PV \rightarrow Pyr)$ | 6 ms |
| Reset potential | -65 mV | $\tau_s(PV \rightarrow PV)$ | 4.3 ms |
| Refractory Period | 2 ms | Pyr cell # | 320 |
| $\tau_s(Pyr \rightarrow Pyr)$ | 2 ms | PV cell # | 80 |

### 2.3. Synapse Model

Excitatory (inhibitory) synaptic events evoke instantaneous jumps (dips) of membrane potentials *V* and their decay over time. The inhibition and excitation decays over different time scales were adopted from an earlier work [30]. The strengths of synaptic connections within assemblies are static over time, whereas the strengths of connections across assemblies are either static or dynamic depending on simulation conditions. When synaptic connections are dynamic, the connections’ strengths follow the spike-time dependent plasticity (STDP) rule summarized in Eq. 2.

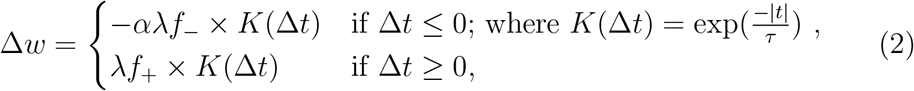

where Δ*t* = *t_post_ − t_pre_* and *f*_+_(*w*) = (1 − *w*)*μ, f_−_*(*w*) = *αμ*. In this study, we used default values included in the NEST package (*α* = 1*, τ* = 20. 0 ms, *μ* = 1. 0*, λ* = 0. 01).

### 2.4. Simulation of local field potentials

Local field potentials (LFPs) reflect population responses. While their origins remain poorly understood, synaptic inputs have been thought to be crucial. Consequently, we simulated LFPs by summing up all (excitatory and inhibitory) synaptic inputs to L2/3 Pyr neurons in PA [31, 32]. The spectral power was calculated by the ‘signal.welch’ routine included in the scipy package [33].

## 3. Results

As shown in Fig. 1, the two assemblies in the low-order area project to the high-order area (HA), and HA projects back to them via nonspecific top-down connections. Initially, the bottom-up connections from both low-order area assemblies are identical and strong enough to drive the HA assembly, when the low-order area assemblies generate sufficiently strong outputs. With these bottom-up connections, when any of the low-order area assemblies become active, HA will fire; that is, the selected low-order assembly and HA will fire together. We assume that as a result, these bottom-up connections will be selectively strengthened via a Hebbian learning rule proposed [34]. Spike-time-dependent-plasticity (STDP) has been suggested as a mechanism that underlies Hebbian learning in the brain [35]. We incorporate STDP in all bottom-up connections in the model and examine the contribution of nonspecific top-down inputs to learning in the bottom-up connections.

### 3.1. Nonspecific top-down signals can sharpen bottom-up connections

In our model, we randomly choose a preferred assembly (PA) to which we introduce 500 Hz external inputs, and a non-preferred assembly (NPA), which receives 100 Hz external inputs, unless stated otherwise; these inputs model sensory inputs from the lateral geniculate nucleus (LGN). The bottom-up connections from PA to HA are expected to grow according to Hebbian learning rule. For the sake of brevity, bottom-up connections from PA and NPA will be referred to as Conn_PA and Conn_NPA, respectively, hereafter. Due to the observations that inter-area connections are layer-specific, all three assemblies consist of superficial (L2/3) and granular (L4) layers, in which excitatory and inhibitory neurons interact with one another via randomly established connections (Table 1). We refer to excitatory neurons as ‘Pyr’ neurons, since most excitatory neurons are shaped like pyramids, and inhibitory neurons as ‘PV’ neurons, since the most common molecular marker of inhibitory neurons is parvalbumin (PV).

We simulate the network for 20 seconds (s) to estimate how Conn_NPA and Conn_PA evolve over time with and without nonspecific top-down inputs onto both NPA and PA (Fig. 1). Fig. 2A, B, and C show the spikes from PA, HA, and NPA for the first 3 s; Pyr and PV neurons are shown in red and blue, respectively. As shown in the figures, Pyr and PV neurons within each assembly fire synchronously several times. We note that the synchronous activity appears first in PA, which receives 500 Hz afferent inputs, and it subsequently appears in L4 neurons of HA and L2/3 neurons of NPA (Fig. 2D). This pattern can be readily explained by the pattern of bottom-up and top-down connections in the model. More importantly, when such sequential activations occur, according to the STDP rule, Conn_PA grow stronger, whereas Conn_NPA grow weaker. Indeed, this sequential activation occurs throughout the simulation. Consequently, Conn_PA grow stronger gradually, but Conn_NPA grow weaker gradually (Fig. 3A), suggesting that bottom-up connections can be selectively strengthened with nonspecific top-down inputs.

**Figure 2:**
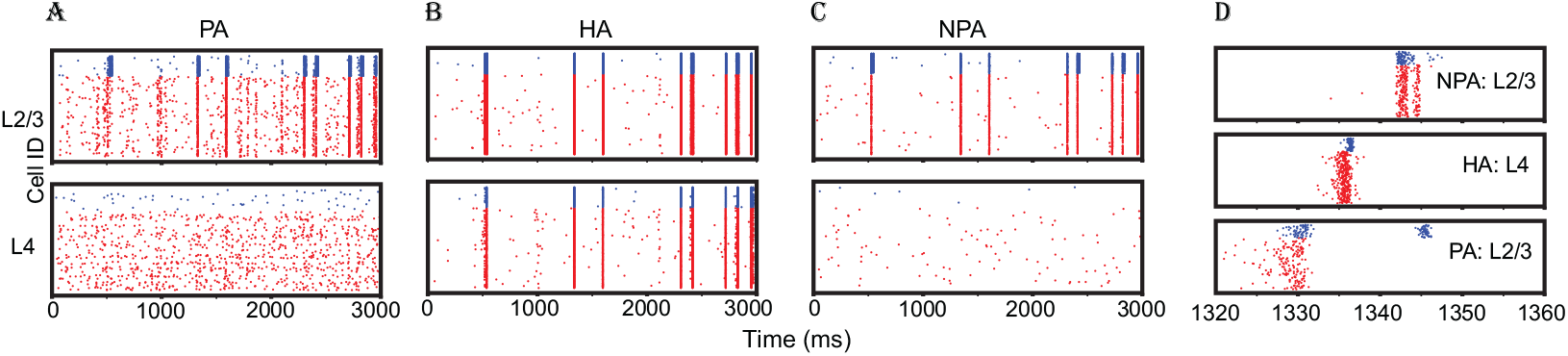
Raster plots of an example simulation. (A), (B), (C), spikes generated in L2/3 and L4 of PA, HA and NPA, respectively. Each dot represents a spike, and spikes from Pyr and PV neurons are shown in red and blue, respectively. For clarity, we show only the first 3 seconds of the simulation. (D), spikes from L2/3 of NPA and PA and those from L4 of HA, between 1320 and 1360 ms.

**Figure 3:**
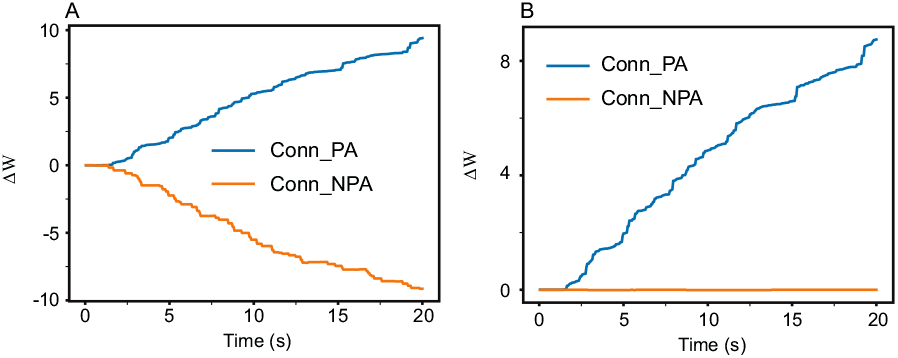
Time course of the change in strength of bottom-up connections. (A), the mean change in weight of bottom-up connections as a function of time. The blue and orange lines represent the mean values for connections from PA to HA and those from NPA to HA, respectively. The mean values are calculated every 5 ms during simulations. (B), the same as (A) but with top-down connections removed.

To further examine the functions of nonspecific top-down inputs, we repeat the simulation without top-down inputs (both to PA and NPA). When top-down connections are removed from the model, Conn_PA increase as before, but Conn_NPA remain unchanged (Fig. 3B). These results suggest that nonspecific top-down connections can reduce the strengths of undesired bottom-up connections (i.e., connections from NPA to HA in this model), keeping the total strength of bottom-up connections at roughly the same level. We further test the effects of top-down inputs on bottom-up connections by varying the strengths of top-down connections to Pyr and PV in L2/3 of PA and NPA. To reduce statistical biases, we conduct 20 simulations, in which networks are independently constructed using the same connectivity rules (see Methods). Fig. 4 shows the total change in bottom-up connection strengths induced during 20-second-long simulations depending on the strengths of top-down connections to PV and Pyr, respectively; the mean and standard deviations from 20 independent simulations (see Methods) are displayed in the figure.

**Figure 4:**
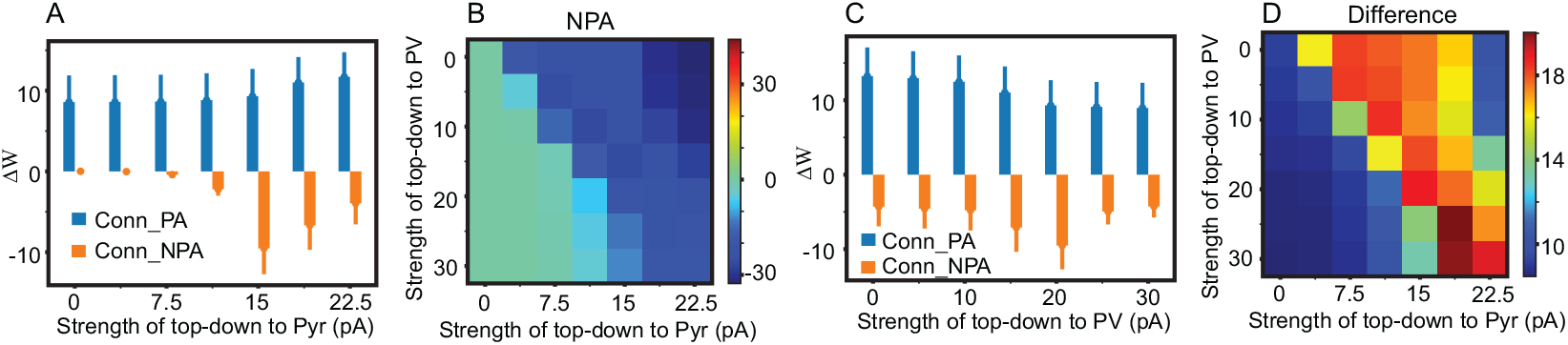
Dependency of the strength of bottom-up connections on the strength of top-down connections. (A), The total change in connection strength of connections from PA and NPA (blue and orange bars, respectively), showing dependence on the strength of top-down inputs to Pyr neurons. Mean values and standard deviations are calculated from 20 independent simulations (see Methods). (B), The total change in connection strength of bottom-up connections from NPA, showing dependence on the strength of top-down connections to both Pyr and PV neurons. (C), The total change in connection strength of connections from PA and NPA, showing dependence on the strength of top-down inputs to PV neurons. (D), The difference in connection strengths of PA and NPA, showing dependence on the strength of top-down connections to both Pyr and PV neurons. The values in all of the figures are calculated using 20 independent 20-second simulations.

We make two observations. First, without top-down inputs to Pyr neurons, the reduction of Conn_NPA is not pronounced (Fig. 4A), confirming that the delayed activity of Pyr neurons in NPA is essential to making Conn_NPA weaker. Second, top-down inputs to PV neurons also have an impact on bottom-up connections’ learning (both Conn_PA and Conn_NPA). As shown in Fig. 4B, the stregnths of Conn_NPA decrease depending on both top-down inputs to Pyr and PV neurons. Additionally, we note that the effect of top-down inputs to PV neurons on Conn_NPA is not monotonic(Fig. 4C). As the strength of top-down connections to PV increases to a threshold of approximately 20 pA, the magnitude of the change in NPA connection weight increases, but after 20 pA, the magnitude decreases.

In contrast, the (positive) change in strength of PA connections monotonically decreases as the strength of top-down connections to PV increases (Fig. 4C). We also note that the difference between Conn_PA and Conn_NPA after 20-second-long simulations is maximal when PV and Pyr neurons receive almost the equivalent amount of top-down inputs (Fig. 4D).

### 3.2. The strength of synchronous oscillatory activity is correlated with the quality of learning

The simulation results above suggest that synchronous activity may be essential in bottom-up connections’ learning. Because the degree of synchronous activity is commonly measured via spectral power of local field potentials (LFPs), we simulate LFPs (Methods) and calculate their spectral power. As shown in Fig. 5A, PA generates oscillatory activity at two frequency bands, 0-20 Hz and 40-80 Hz. To assess how much synchronous oscillatory activity contributes to bottom-up connections’ learning, we quantify the changes in strengths of bottom-up connections (Conn_PA and Conn_NPA) and the total power of the two frequency bands (0-20 and 40-80 Hz) for all 20 simulations. Fig. 5B shows that the magnitudes of changes in bottom-up connections are positively correlated with the oscillatory power in PA, supporting that cortical rhythms (i.e., synchronous oscillatory neural activity) drive bottom-up connections’ learning.

**Figure 5:**
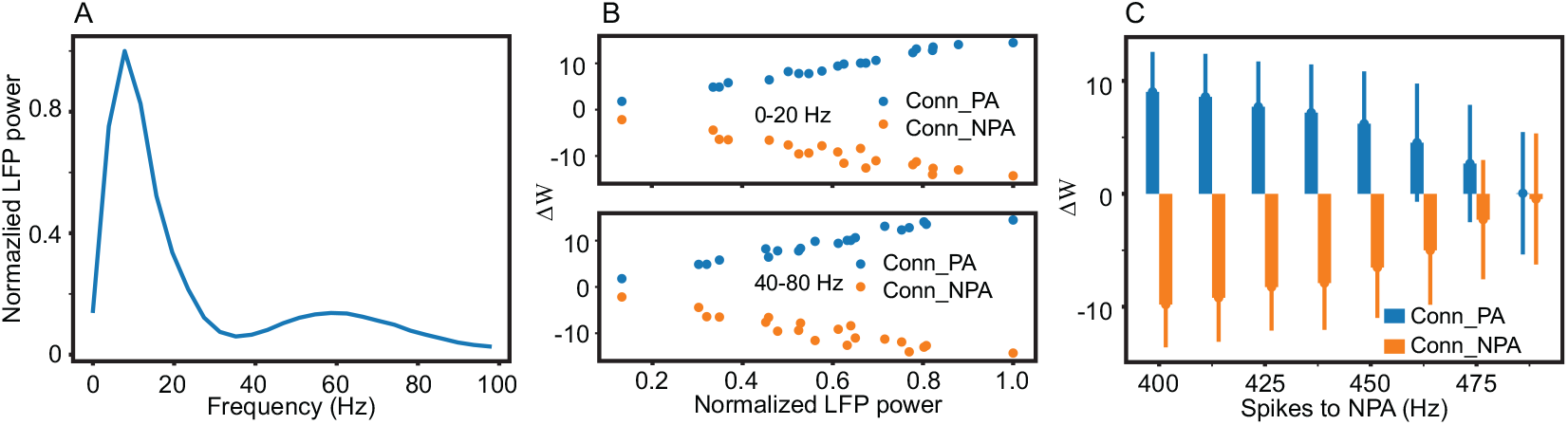
Contribution of synchronous oscillatory activity to learning and the robustness of the learning process. (A), Spectral power of LFPs in PA, normalized to the maximum value. (B), Correlations between total change in connection strength and spectral power of LFPs in PA in two frequency bands (0-20 and 40-80 Hz). For each simulation (out of 20), we quantify the spectral power and total change in bottom-up connection strength. Spectral power in each frequency band is normalized to the maximum power obtained during 20 simulations. Blue and orange dots represent connections originating from PA and NPA, respectively. (C), Total change in connection strength, showing dependence on the frequency of external inputs to NPA; the default inputs to PA and NPA are 500 and 100 Hz.

In the model, if NPA, instead of PA, generates synchronous outputs, Conn_NPA, instead of Conn_PA, become stronger. If the inputs to NPA are not strong enough, the growth of undesired connections (from NPA to HA) are negligible, on average. Then, how sensitive is this learning mechanism to the inputs to NPA? To answer this question, we increase external inputs to NPA up to 475 Hz. Fig. 5C shows that the growth of Conn_PA decreases as the frequency of inputs to NPA increases. But importantly, despite the reduction in magnitude, the Conn_PA grow stronger, whereas Conn_NPA grow weaker, on average, even when the external inputs to NPA are 90% of those to PA. This result suggests that nonspecific top-down connections can be useful to selectively strengthen the connections even when all low-order assemblies receive relatively similar afferent inputs.

### 3.3. Top-down connections can also evolve in a target-specific way via learning

So far, we have assumed that only bottom-up connections have plasticity, but top-down connections can also have plasticity. That is, top-down connections can evolve over time. Naturally, two questions arise. First, will bottom-up connections change the same way when top-down connections are given plasticity? Second, how do top-down connections change? To answer these questions, we give top-down connections STDP mechanisms and repeat the same experiments. In simulation, we impose the maximum strength of top-down connections to be 30 pA; without such limitation, the system can show run-away excitation due to strong positive feedback loops established between PA (NPA) and HA. We note (1) that bottom-up connections change in the same ways as they do with stationary top-down connections (Fig. 6A and B) and (2) that top-down connections change depending on their targets (Fig. 6C and D). Specifically, top-down connections targeting NPA grow stronger, while top-down connections targeting PA grow weaker, consistent with experimental findings [36].

**Figure 6:**
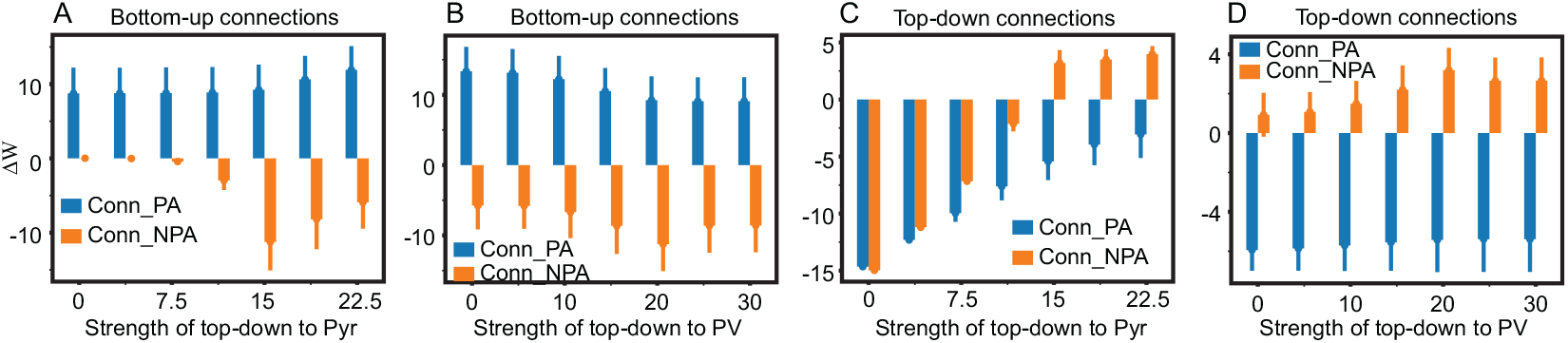
Simulation results with dynamically evolving top-down connections. (A), Change in bottom connection strength induced during long simulations, showing dependence on initial strength of top-down connections to Pyr. Blue and orange colors represent bottom-up connections originating from PA and NPA, respectively. (B), the same as (A), but showing dependence on initial strength of top-down connections to PV. (C), Change in top-down connection strengths induced during simulations, showing dependence on initial strength of top-down connections to Pyr. Blue and orange colors represent top-down connections targeting PA and NPA, respectively. (D), the same as (C), but showing dependence on initial strength of top-down connections to PV.

### 3.4. Learning of STDP connections can be dynamically turned on or off

Neural networks operate in two different modes, ‘training’ and ‘inference’. In the inference mode, weights are frozen, and actual functions are performed by neural networks. The switching between the two modes is fundamental to the neural networks’ operations, raising the possibility that the brain can turn on or off learning depending on the context. We note that recent experimental studies [37, 38] suggest that vasoactive intestinal peptide (VIP)-expressing inhibitory neurons play important roles in the brain’s learning. Together with the observation [39] that somatostatin (SST)-expressing inhibitory neurons and VIP neurons mutually inhibit each other, we hypothesize that the interplay between SST and VIP neurons regulates the learning of bottom-up connections from PA and NPA to HA. To address this hypothesis, we incorporate SST and VIP neurons into the model. Based on experimental observations [39, 40], we set SST neurons to inhibit all other neuron types except themselves and VIP neurons to inhibit SST neurons exclusively (Fig. 7A); see Table 3 for model connectivity.

**Table 3:**
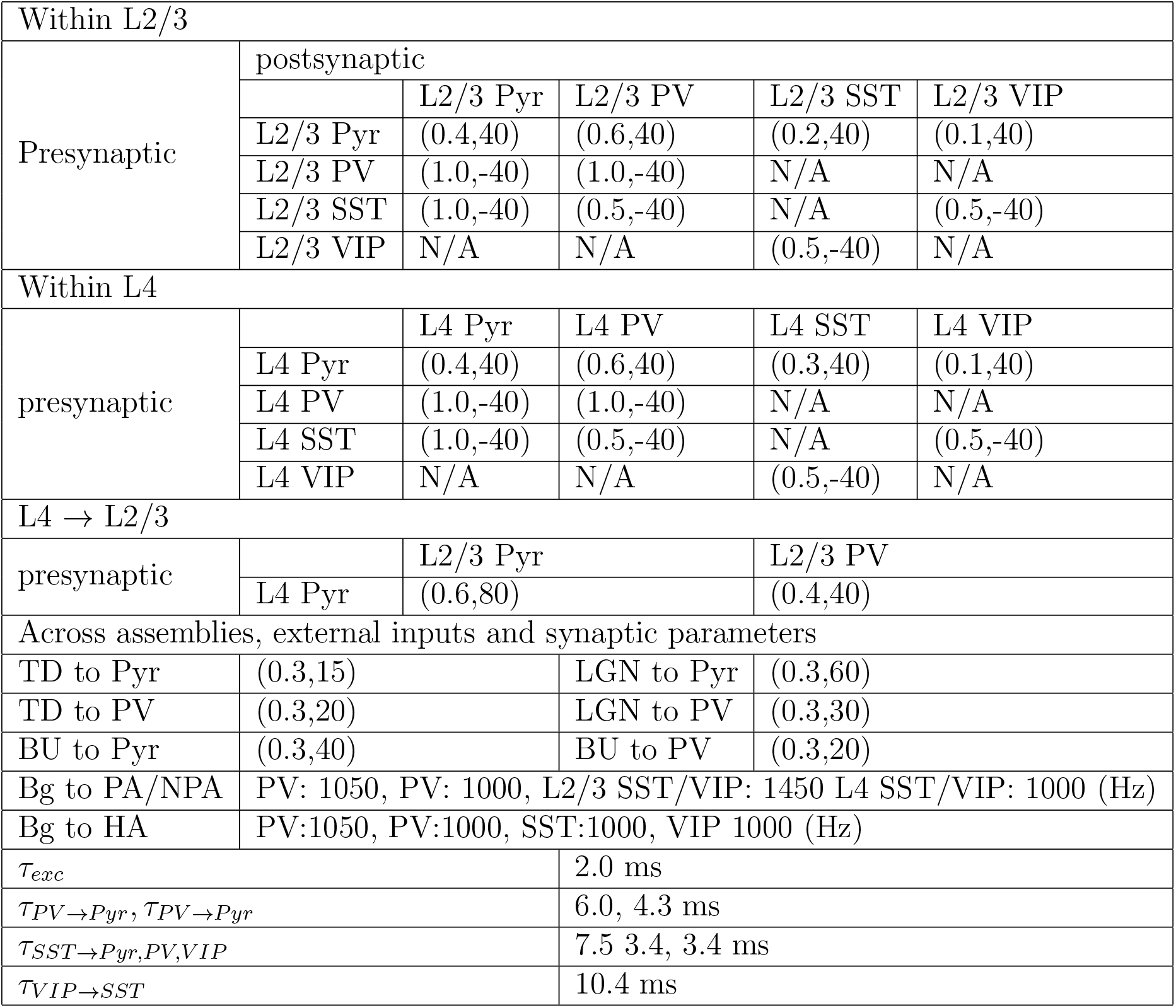
Connections in the network model consisting of Pyr and three inhibitory neurons. Below, the connection probability and strength of each connection type is shown in parentheses; the unit for connection strength is pA. TD, BU, LGN and Bg represent top-down, bottom-up, LGN inputs and Background inputs, respectively. The external inputs to neurons (Bg) deliver 100 pA at their peaks.

|  |  |  |  |  |  |
| --- | --- | --- | --- | --- | --- |
| Within L2/3 |  |  |  |  |  |
| Presynaptic | postsynaptic |  |  |  |  |
|  |  | L2/3 Pyr | L2/3 PV | L2/3 SST | L2/3 VIP |
|  | L2/3 Pyr | (0.4,40) | (0.6,40) | (0.2,40) | (0.1,40) |
|  | L2/3 PV | (1.0,-40) | (1.0,-40) | N/A | N/A |
|  | L2/3 SST | (1.0,-40) | (0.5,-40) | N/A | (0.5,-40) |
|  | L2/3 VIP | N/A | N/A | (0.5,-40) | N/A |
| Within L4 |  |  |  |  |  |
| presynaptic |  | L4 Pyr | L4 PV | L4 SST | L4 VIP |
|  | L4 Pyr | (0.4,40) | (0.6,40) | (0.3,40) | (0.1,40) |
|  | L4 PV | (1.0,-40) | (1.0,-40) | N/A | N/A |
|  | L4 SST | (1.0,-40) | (0.5,-40) | N/A | (0.5,-40) |
|  | L4 VIP | N/A | N/A | (0.5,-40) | N/A |
| L4 → L2/3 | | | | | |
| presynaptic |  | L2/3 Pyr |  | L2/3 PV |  |
|  | L4 Pyr | (0.6,80) |  | (0.4,40) |  |
| Across assemblies, external inputs and synaptic parameters |  |  |  |  |  |
| TD to Pyr | (0.3,15) |  | LGN to Pyr | (0.3,60) |  |
| TD to PV | (0.3,20) |  | LGN to PV | (0.3,30) |  |
| BU to Pyr | (0.3,40) |  | BU to PV | (0.3,20) |  |
| Bg to PA/NPA | PV: 1050, PV: 1000, L2/3 SST/VIP: 1450 L4 SST/VIP: 1000 (Hz) |  |  |  |  |
| Bg to HA | PV:1050, PV:1000, SST:1000, VIP 1000 (Hz) |  |  |  |  |
| $\tau_{exc}$ | | | 2.0 ms | | |
| $\tau_{PV \rightarrow Pyr}, \tau_{PV \rightarrow Pyr}$ | | | 6.0, 4.3 ms | | |
| $\tau_{SST \rightarrow Pyr, PV, VIP}$ | | | 7.5 3.4, 3.4 ms | | |
| $\tau_{VIP \rightarrow SST}$ | | | 10.4 ms | | |

**Figure 7:**
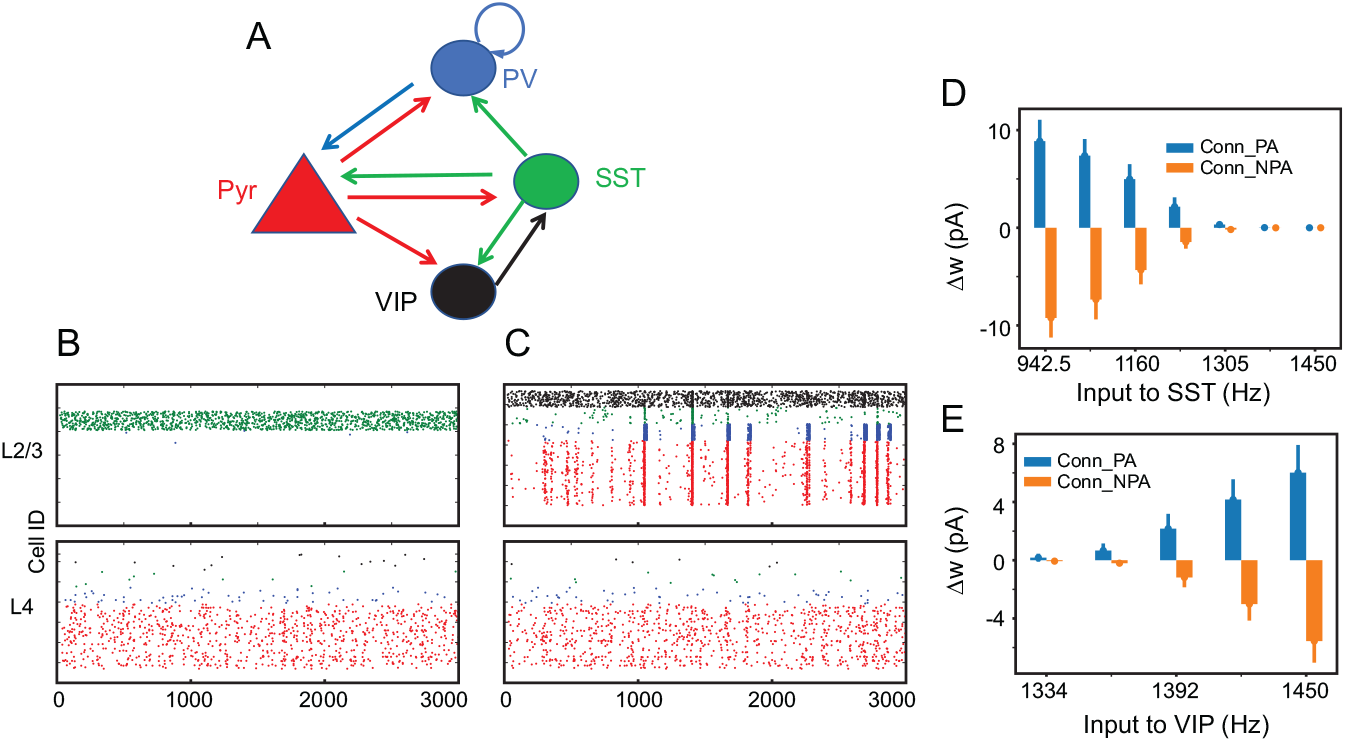
Simulation results with three inhibitory neuron types. (A), Connectivity among Pyr and three inhibitory neuron types; see Table 3 for details. All neuron types have identical sub-threshold dynamics (Table 2). (B), An example of simulation without external inputs to VIP neurons. (C), the same as (B), but with external inputs (1450 Hz) to VIP neurons. (D), Δ*W* of Conn_PA and Conn_NPA with varying external inputs to SST neurons. The *x*-axis represents the frequency of external inputs to SST neurons. In these simulations, no external inputs are introduced to VIP neurons. (E), Δ*W* with varying external inputs to VIP neurons; in all these simulations shown in (E), SST neurons always receive 1450 Hz external inputs.

In our simulations, when SST neurons receive sufficient external (1450 Hz) inputs and became active, Pyr neurons become quiescent and do not fire synchronously (Fig. 7B). In contrast, when VIP cells are sufficiently depolarized by the external (1450 Hz) inputs, SST activity is suppressed, and Pyr neurons fire synchronously (Fig. 7C). As Pyr synchronous firing is essential in STDP connections’ learning in the model, VIP cell activity could play an important role in the bottom-up connections’ learning. To test this possibility, we vary external inputs to SST and VIP neurons and quantified the changes in strengths of Conn_PA and Conn_NPA. As the frequency of inputs to SST neurons increase, the changes in the strengths of Conn_PA and Conn_NPA become smaller (Fig. 7D). While maintaining 1450 Hz inputs to SST neurons, we gradually increase the frequency of inputs to VIP neurons. The changes in the strengths of Conn_PA and Conn_NPA are positively correlated with the frequency of inputs to VIP neurons (Fig. 7E). These results support that the interplay between VIP and SST cells can dynamically regulate learning in the brain.

### 3.5. Brain areas can be independently trained even with shared top-down connections

Our simulation results suggest that nonspecific connections can selectively enhance connections from one brain area to another. If one assembly in a high-order area (HA) is exclusively connected to a single low-order area, non-specific top-down connections can exclusively sharpen the desired bottom-up connections without perturbing the bottom-up connections from other low-order areas. However, it is well known that brain areas are densely interconnected and that individual neurons receive inputs from multiple areas; see [9] for a review. Thus, we cannot exclude the possibility that a single HA is reciprocally connected with multiple low-order areas. If so, the activity in a low-order area can change the bottom-up connections from other low-order areas to the HA, which can create undesired crosstalk between two low-order areas. For instance, if a HA integrates auditory and visual inputs, continuous auditory inputs will make the HA insensitive to visual inputs, which can be harmful. Thus, it seems natural to assume that the brain can selectively turn on learning in individual brain areas. Then, how does the brain control individual areas’ learning?

Inspired by our simulation results (Fig. 7), we hypothesize that afferent inputs to VIP neurons can serve as control signals for learning. To investigate this hypothesis, we construct a model in which two identical areas (areas 1 and 2) are connected to the same HA. As shown in Fig. 8A, areas 1 and 2 are identical in terms of the structure and external inputs, with each containing a PA and an NPA. In each assembly, Pyr neurons interact with PV, SST and VIP neurons (Table 3), and we introduce afferent inputs (control signals, 1450 Hz) to VIP in either areas 1 or 2, while maintaining the 1450 Hz inputs to L2/3 SST neurons. As SST neurons are known to mediate lateral inhibition via di-synaptic connections [39], we connect L2/3 Pyr neurons in one assembly to L2/3 SST neurons in another. Similarly, the lateral excitatory connections are also implemented by connecting L2/3 Pyr neurons across assemblies. The connection probability and strength of both lateral connections are 0.05 and 10 pA. It should be noted that these lateral connections exist only within the same area.

**Figure 8:**
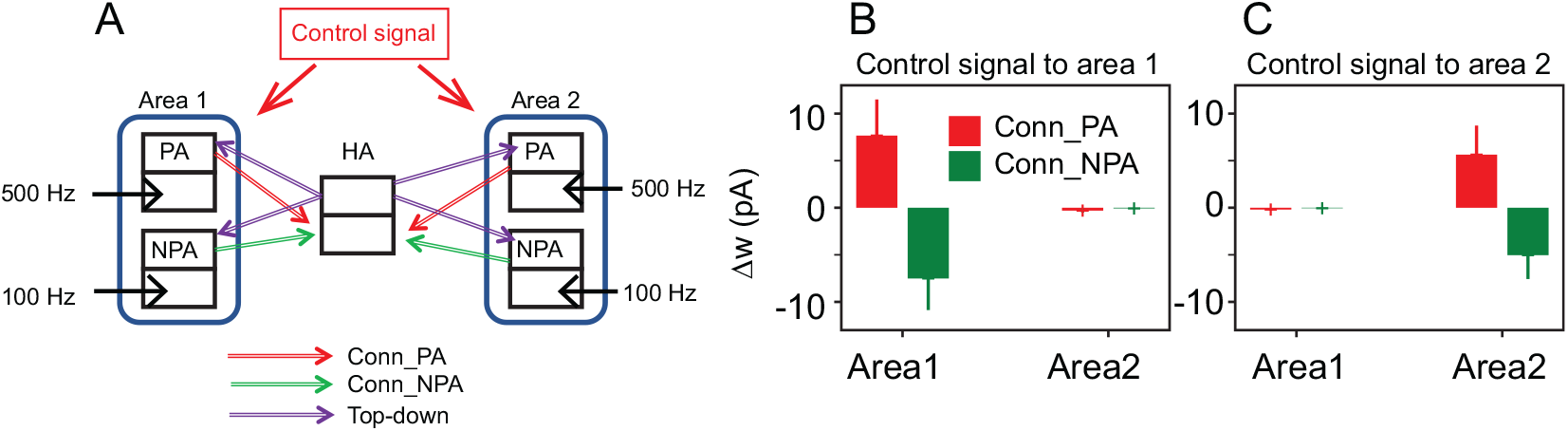
Simulation results with two low-order areas. (A), The schematics of the model. Each assembly consists of Pyr, PV, SST and VIP neurons. The assemblies are identical to those used to generate the results in Fig. 7. (B), Δ*W* of areas 1 and 2 when the control signals are projected to area1 only. (B), Δ*W* of areas 1 when the control signals are projected to area2 only. The mean values and standard deviations are estimated from 20 independent simulations, each of which lasted 20 seconds.

Fig. 8B shows the mean values and standard deviations of 20 independent simulations with control signals given to either area 1 or area 2. When area 1 VIP neurons receive control signals (1450 Hz), only connections (Conn_PA and Conn_NPA) from area 1 to HA change. Similarly, when control signals are introduced to area 2, only bottom-up connections from area 2 change. These results support that the learning of STDP connections can be dynamically regulated via inputs to VIP neurons.

### 3.6 Simulation results are not sensitive to neuron models

First, a different variation of current-based LIF neurons is used. In our reference model discussed above, synaptic currents caused instantaneous jumps in membrane potentials. These instantaneous jumps in membrane potentials do not capture the gradual increase in membrane potentials of biological neurons evoked by spikes. Thus, we set LIF neurons to have an ‘alpha postsynaptic function’ (Eq. 3) that generates gradual increases in membrane potentials.

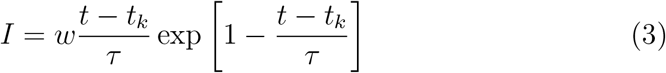

where *w* is the strength of weights and *t_k_* and *τ* are a spike time and a time constant for synaptic inputs, respectively. As LIF neurons with alpha postsynaptic functions have different response properties, we modify the parameters of the model, which are listed in Table 4. After the modifications, we measure the changes in strengths of Conn_PA and Conn_NPA from 20 independent simulations and found the results to be equivalent to those described above (Fig. 9A).

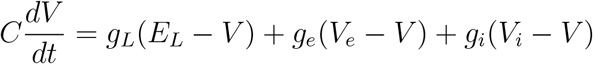

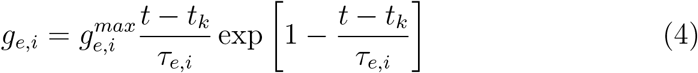

where *g_e_, g_i_, g_L_* represent the conductances of excitatory synaptic inputs, inhibitory synaptic inputs, and leak currents, respectively; *V_e_, V_i_, E_L_* represent the reversal potentials for excitatory synaptic inputs, inhibitory synaptic inputs, and leak currents, respectively; *τ_e,i_* represent the time constants for synaptic inputs; *C* is the capacitance of the neuron; and 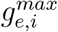 correspond to the synaptic weights in Table 5. All parameters are taken from the default values of the NEST native implementations (‘iaf_psc_alpha’ and ‘iaf_cond_alpha’).

These results support that our findings are not contingent upon the selection of neuron models.

**Table 4:** Connections in the network model with alternative LIF neurons, which have ‘alpha’ postsynaptic function. Below, the connection probability and strength of each connection type is shown in parentheses; the unit of connection strength is pA. TD, BU, LGN, and Bg represent top-down, bottom-up, LGN inputs, and Background inputs, respectively. The external inputs to neurons (Bg) deliver 100 pA at their peaks. Except *α*, all parameters for STDP connections are the same as those used above.

| Presynaptic | Postsynaptic |  |  |  |  |
| --- | --- | --- | --- | --- | --- |
|  |  | L2/3 Pyr | L2/3 PV | L4 Pyr | L4 PV |
|  | L2/3 Pyr | (0.4,40) | (0.6,40) | N/A | N/A |
|  | L2/3 PV | (1.0,-40) | (1.0,-40) | N/A | N/A |
|  | L4 Pyr | (0.6,40) | (0.4,40) | (0.4,40) | (0.6,40) |
|  | L4 PV | N/A | N/A | (1.0,-40) | (1.0,-40) |
| Across Assemblies and external inputs |  |  |  |  |  |
| TD to Pyr | (0.3,15) |  | LGN to Pyr | (0.3,20) |  |
| TD to PV | (0.3,20) |  | LGN to PV | (0.3,30) |  |
| BU to Pyr | (0.3,40) | | $\tau_{exc}$ | 1.0 ms | |
| BU to PV | (0.3,40) | | $\tau_{PV \rightarrow Pyr}$ | 3.0 ms | |
| Bg to Pyr | 1050 Hz | | $\tau_{PV \rightarrow PV}$ | 2.1 ms | |
| Bg to PV | 1100 Hz | | STDP param | $\alpha = 1.5$ | |

**Table 5:** Connections in the network model with conductance-based LIF neurons. Below, the connection probability and strength of each connection type are shown in parentheses; the unit of connection strengths is nS. TD, BU, LGN, and Bg represent top-down, bottom-up, LGN inputs, and Background inputs, respectively. The external inputs to neurons (Bg) deliver 15 nS at their peaks. LA refers to both PA and NPA. Except *α*, all parameters for STDP connections are the same as those used above.

|  |  |  |  |  |  |
| --- | --- | --- | --- | --- | --- |
| Presynaptic | Postsynaptic |  |  |  |  |
|  |  | L2/3 Pyr | L2/3 PV | L4 Pyr | L4 PV |
|  | L2/3 Pyr | (0.05,2) | (0.5,1.0) | N/A | N/A |
|  | L2/3 PV | (0.8,-12) | (0.5,-8) | N/A | N/A |
|  | L4 Pyr | LA: (0.4,6)<br>HA: (0.4,2) | (0,2,2) | (0.05,2) | (0.5,10) |
|  | L4 PV | N/A | N/A | LA: (0.8,-12)<br>HA: (0.8,-12) | LA: (0.5,-8)<br>HA: (0.5,-2) |
| Across Assemblies and external inputs |  |  |  |  |  |
| TD to Pyr | (0.05,3) |  | LGN to Pyr | (0.3,20) |  |
| TD to PV | (0.05,4) |  | LGN to PV | (0.3,10) |  |
| BU to Pyr | (0.08,25) | | $\tau_{exc}$ | 1.0 ms | |
| BU to PV | (0.08,25) | | $\tau_{PV \rightarrow Pyr}$ | 3.0 ms | |
| Bg to Pyr | 25 Hz | | $\tau_{PV \rightarrow PV}$ | 2.1 ms | |
| Bg to PV | 25 Hz | | STDP param | $\alpha = 3.0$ | |

**Figure 9:**
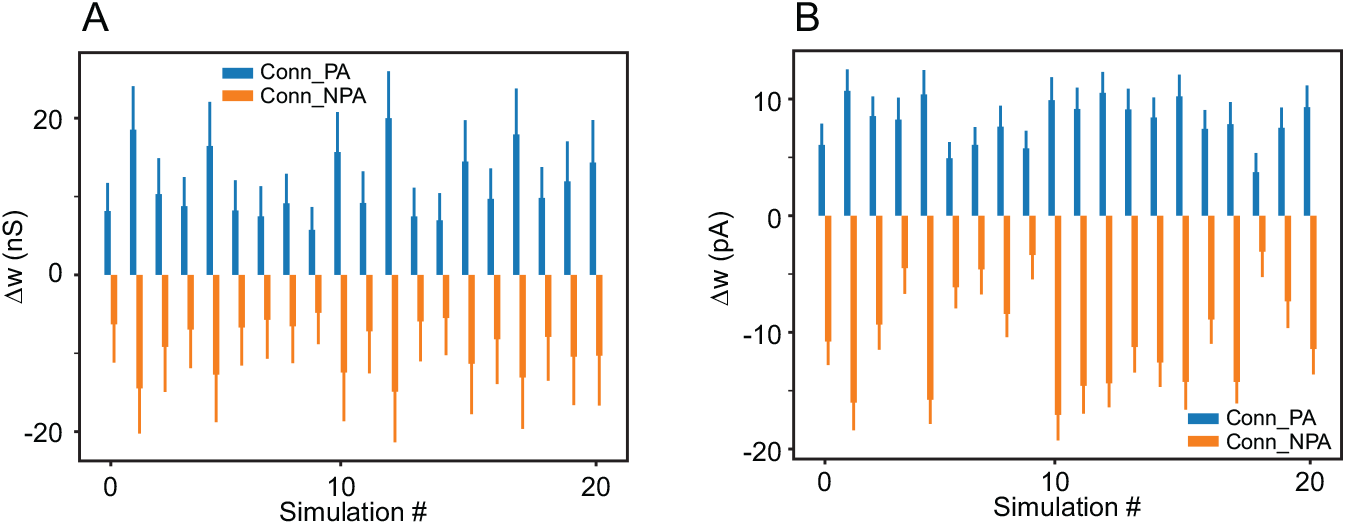
Simulation results with alternative neuron models (A) Δ*W* of Conn_PA and Conn_NPA using current-based LIF neurons with alpha-function. We display the mean values and standard deviations of Δ*W* in all 20 simulations, each of which lasted 20 seconds. (B), Δ*W* of Conn_PA and Conn_NPA with conductance-based LIF neurons with alpha-function. We display the mean values and standard deviations of Δ*W* in all 20 simulations, each of which lasted 50 seconds.

## 4. Discussion

In this section, we describe the implications of our simulation results in more detail.

### 4.1. Implications for learning of Spiking Neural Networks

Spiking neural networks (SNNs) are the building blocks of the brain and are known to be power-efficient. With the brain’s general intelligence in mind, it seems only natural to construct artificial SNNs to advance ML; see [41] for a review. However, training SNNs can be challenging, and their exact operating principles remain elusive; see [42] for a review. In principle, an understanding of the mechanisms by which the brain learns could be used to develop learning algorithms for SNNs. STDP has been thought to underlie, in part, the brain’s learning capabilities [43]. However, STDP (in general, variations of Hebbian plasticity) are known to drive SNNs into highly active states [28]. This causes the neurons to become insensitive to external inputs and the synaptic weights to increase continuously, rendering SNNs useless. Zenke et al. [28] propose that the homeostatic process, which regulates the growth of synaptic strengths, is necessary to block SNNs from moving into a persistent state of high activity. As nonspecific top-down inputs can suppress undesired connections effectively, we propose that experimentally-observed nonspecific top-down connections can serve as homeostatic processes critical for STDP-based learning of SNNs.

### 4.2. Functions of top-down inputs to PV neurons

In the model, nonspecific top-down inputs to Pyr neurons can reduce undesired bottom-up connections from NPA by inducing delayed activity in NPA. Additionally, the results in Fig. 4 suggest that top-down inputs to PV neurons also contribute to bottom-up connections’ learning. How do PV neurons contribute to this learning process?

We propose that PV neurons prevent Pyr neurons from firing in a way that disrupts the temporal order of spiking across assemblies, thus allowing for the strengthening of PA bottom-up connections and the weakening of NPA bottom-up connections. For example, PV neurons may prevent L2/3 Pyr neurons in PA from firing continuously. If L2/3 Pyr neurons in PA fired continuously when driving HA, they could fire either before or after the firing of HA neurons and thus by STDP mechanisms, Conn_PA could be either strengthened or weakened. Similarly, if L2/3 Pyr neurons of NPA fired continuously after the firing of HA neurons, they could potentially fire just before the firing of L4 Pyr neurons of NPA, leading to an increase in the strength of Conn_NPA by STDP mechanisms. The PV neurons may prevent these phenomena, thus allowing STDP to produce an increase in Conn_PA and a decrease in Conn_NPA, as observed.

We note that PV neurons in PA and NPA have different responses to local and HA inputs (Fig, 2D). In PA, PV neurons (not Pyr neurons) respond to top-down inputs. Since PV neurons in PA fire in response to top-down inputs, Pyr neurons receive inhibition which prevents them from firing in response to top-down inputs. This minimizes the chance of the reduction of Conn_PA. In NPA, on the other hand, PV neurons respond mostly to local inputs from NPA, not top-down inputs. That is, PV neurons in NPA provides negative feedback inputs to Pyr neurons in NPA, making the active period of NPA Pyr neurons to be brief. Interestingly, the undesired firing of Pyr neurons occurs more frequently when top-down inputs are strong. Consequently, top-down inputs to PV should be proportionally increased to ensure the proper learning of bottom-up connections; this is consistent with the balanced top-down inputs to Pyr and PV neurons observed experimentally [44].

### 4.3. The function of synchronous activity in interareal connections

The communication-through-coherence (CTC) theory proposes that cortical rhythms (synchronous and oscillatory neural activity) subserve interareal interactions by increasing the efficacy of afferent inputs to target areas [26]. Consistent with this theory, synchronous activity in our model reliably propagates across assemblies via stochastic connections. More importantly, we note that synchronous activity in our model not only enhances the efficacy of interareal communications, but also contributes to interareal connections’ learning in two ways. First, when synchronous activity is generated and propagates among areas, postsynaptic neurons in the target area fire reliably after presynaptic neurons fire. This ensures the temporal gaps between pre- and postsynaptic neurons, allowing the synaptic weights connecting them to change robustly. Second, the induced inhibition during synchronous firing prevents Pyr from incorrectly firing, which makes the learning process more reliable. Thus, our simulation results propose a possible extension of CTC theory [26].

### 4.4. Potential coordination between specific and nonspecific top-down inputs

We find that two conditions need to be satisfied for reliable learning of bottom-up connections. First, a selected assembly should be active (or activated) more than other assemblies. Second, the most active cell assembly needs to generate synchronous activity. Conversely, the proposed learning process will not be effective when multiple assemblies receive almost the equivalent amount of afferent inputs or when the desired assembly (i.e., PA in the model) does not generate synchronous activity. However, these limits may not be fundamental and may be dynamically regulated by selective attention, which is known to be capable of biasing competition and amplifying synchronous activity. If selective attention is directed to NPA, the synchronous activity of NPA is amplified, and Conn_NPA, rather than Conn_PA, become stronger. In contrast, the growth of Conn_PA accelerates if selective attention is directed to PA. Interestingly, an earlier study proposed that top-down inputs impinging onto deep layers are target-specific [45]. Thus, selective attention may regulate this learning process via stimulation of deep layers. In the future, we plan to address this possibility by extending our model to incorporate deep layers and inhibitory neuron types known to be associated with interlayer interactions.

## 5. Conclusion

The brain is a group of recurrent networks, in which high and low order areas interact reciprocally with one another. It is generally accepted that bottom-up processing is central to generating perception as demonstrated in deep learning [7]. However, the functions of top-down inputs remain poorly understood. Since most theories assume that top-down inputs are target specific, they cannot explain the functions of experimentally reported nonspecific top-down inputs [21, 20, 22, 23, 24]. Here, we employ network models to investigate how nonspecific top-down connections contribute to the dynamic organization of bottom-up connections. Our simulation results propose 1) that nonspecific top-down inputs mediate temporal information to low-order areas to induce the specificity in bottom-up connections and 2) that cortical rhythms are essential in this learning process.

We will extend this study in two different directions. First, we will further study how the coordination between superficial and deep layers can contribute to the brain’s learning. Second, we will examine if nonspecific top-down connections can help SNNs learn practical tasks.

## Acknowledgements

We thank the Allen Institute founder, Paul G. Allen, for his vision, en-couragement, and support.

